# THE PALISADE LAYER OF THE POXVIRUS CORE IS COMPOSED OF FLEXIBLE A10-TRIMERS

**DOI:** 10.1101/2023.05.24.542031

**Authors:** Jiasui Liu, Simon Corroyer-Dulmont, Vojtěch Pražák, Iskander Khusainov, Karola Bahrami, Sonja Welsch, Daven Vasishtan, Agnieszka Obarska-Kosińska, Sigurdur R. Thorkelsson, Kay Grunewald, Emmanuelle R. J. Quemin, Beata Turoňová, Jacomina Krijnse Locker

## Abstract

Although vaccinia virus (VACV) is the best studied poxvirus, the structure of the mature virus (MV) remains poorly understood. Its asymmetric shape, size and compactness poses a major challenge for electron microscopy (EM) analysis, including cryoEM. Sub-viral particles, in particular membrane-free viral cores, may overcome these limitations. We compare cores obtained by detergent-stripping MVs with cores in the cellular cytoplasm, early in infection. By combining cryo-electron tomography (cryoET), subtomogram averaging (STA) and AlphaFold2 (AF2), abundant core-structures are analyzed, focusing on the prominent palisade layer on the core surface. On detergent-stripped cores, the palisade is composed of densely packed trimers of the major core protein A10. On the core surface they display a random order and their classification indicate structural flexibility. On cytoplasmic cores A10 is organized in a similar manner, indicating that the structures obtained *in vitro* are physiologically relevant. CryoET and STA also uncover unexpected details of the layers beneath the palisade both on *in vitro* and *in situ* cores, that are compared to AF2 structure predictions of known VACV core-associated proteins. Altogether, our data identify for the first time the structure and molecular composition of the palisade units. The results are discussed in the context of the VACV replicative cycle, the assembly and disassembly of the infectious MV.

## Introduction

The recent emergence of monkeypox virus infections in Europe and North America has rekindled an interest in poxviruses. The prototype of the poxviruses, vaccinia virus (VACV), was used previously as a live vaccine to eradicate smallpox and has been studied extensively, biochemically, genetically, and morphologically (reviewed in by B. Moss [1]). However, despite extensive investigation, questions remain about the biogenesis and structure of the mature virion (MV)[2].

The infectious mature virion (MV) is a quasi-brick shaped particle measuring roughly 250×350×200 nm. It is composed of an oval core enclosing the viral genome as well as the machinery required for transcription early in infection. This core is surrounded by a lipid bilayer acquired during assembly; two so-called lateral bodies are located on the elongated sides of the core-brick, underneath the viral membrane [3]. The MV is composed of roughly 200 proteins. Treatment of MVs with a non-ionic detergent and a reducing-agent results in a membrane-free, stable, core [4], which is widely used to discriminate viral proteins associated with the membrane and the core. Thus, the membrane-free core is known to be composed of at least five abundant proteins; gene products of A3, A4, A10, L4 and F17 [2]. With the exception of F17, these proteins are also associated with the incoming, cytoplasmic core, early in infection [5].

Although the MV has been studied extensively by electron microscopy (EM), including cryoEM, its precise structure remains elusive. Structural analyses are hampered in particular by the compactness of the virion and its asymmetric shape. Thus, sub-viral structures have also been studied, in particular the membrane-free and detergent-stripped core, by classical negative staining EM [6], [7] and by cryoEM [8]. Such cores display a dense spike-like layer, the so-called palisade, on their surface. The cryoEM data by the team of Jacques Dubochet proposed that the palisade units are hollow tubes, roughly 20nm long and 10nm in diameter, which are tightly packed on the core surface in a disordered and ordered way, forming a hexagonal lattice with a 9.7 nm spacing. Analyzing the intact MV by cryo-electron tomography (cryoET), Cyrklaff et al. concluded that the palisade-units on the core surface were 8nm in length and 5nm in diameter and arranged in hexagonal lattices [3]. A recent publication studying VACV assembly by cryoET in the periphery of infected cells, also proposed an ordered arrangement of the palisade on fully assembled virions. The model in this study displayed the palisade as trimeric pillars with projecting lobes that mediate interpillar contacts [9].

Although the major proteins associated with membrane-free cores are known, it is unclear which of these proteins make up its prominent features, such as the core wall and the characteristic palisade[2]. Wilton et al., using antibody labeling, suggested that the palisade is made up by the gene product of A3 [7].

In the present study we apply cryoET and subtomogram averaging (STA) to study the structure and molecular composition of the palisade both *in vitro* and *in situ.* From the *in vitro* data we obtain a structure of the palisade units at less than 7.7Å resolution. The latter faithfully represents the structure of the palisade *in situ*, on the surface of the incoming core early in infection. We apply AlphaFold2 [10] to predict the 3D structure of the five major core proteins; by flexible fitting of the predicted structure of the gene product of A10 into the obtained map, we show that the palisade units are made up of trimers of this core protein. The relevance of these data, including additional features we observe in our cryo-tomograms, are discussed with respect to the biology of poxvirus infection.

## Results

### Sample preparation of Vaccinia virus cores for cryo electron tomography

For structural analysis of the viral core of Vaccinia virus by cryo electron tomography (cryoET), we first optimized sample preparation to obtain intact cores *in vitro*. Based on previous studies, short incubation with 0.1% NP-40 and 10 mM freshly prepared dithiothreitol (DTT) diluted in 50 mM Tris-Cl pH 8.5 appeared to be the optimal conditions. These freshly prepared cores are structurally intact and remain associated with lateral bodies, while storing them at −80°C results in a considerable number of broken cores. Sonication prior to vitrification is essential to obtain a sufficient number of individual cores on cryoEM grids. Vitrified samples are inspected in a Titan Krios G4 (Thermo Scientific) TEM and areas containing single cores located in the holes of the grids selected for cryoET. In total, 34 tilt series were acquired and reconstructed into tomograms, yielding 50 cores that were used for further analysis. For comparison, supplemental Figure S1 and Movie S1 display a typical MV from purified preparation, exemplifying the complexity of its structure.

### The cores of Vaccinia virus prepared *in vitro* display three layers

Reconstructed tomograms of the cores prepared *in vitro* confirm the overall organization observed previously (Fig.1, Movie S2) [8]. Upon removal of the viral membrane the core acquires a brick-like shape with average dimensions of 200 nm by 320 nm by 200 nm. The inside of the core is sparsely filled with electron-densities possibly representing the viral genome in complex with DNA-binding proteins or the transcription machinery packaged during assembly [11]. It is enclosed by at least three layers (Fig. 1b). The most prominent is the outer palisade layer composed of protrusions with a tube-like shape of ~11 nm in height and diameter of 8 nm (Fig. 1b). Being subunits of the palisade, we will refer to these tubular structures as stakes throughout this study. The stakes do not form a regular lattice (Fig. 1b, d) in contrast to what has been described previously [8], [9], [12]. At their top, the stakes are often connected to their neighbors with thin threats (Fig. 1d). The ring structure reported in [9] is integrated into the palisade (Fig 1c). Below the palisade is the middle layer, the inner core wall with a thickness of 4-5 nm (Fig. 1e). From the side, the layer resembles a lipid bilayer (Fig. 1e), however it appears as stripes when viewed from the top with very weak perpendicular connections between the stripes. Interestingly, the arrangements of the stakes in the palisade layer and the stripes in the inner core wall layer do not seem to be aligned (Movie S2). The most inner layer is roughly 3 nm below the inner core wall and has a thickness of less than 2 nm. The content of this layer appears irregular within each core: it can contain 1 or several rows (Fig. 1b, f) and is sometimes even absent. Strong densities are occasionally found to be connected to the top of the stakes (Fig. 1d). Consistent with previous data and the gentle sample preparation procedure we used, these *in vitro* cores also contain two large blob-like structures on both sides: the lateral bodies. Our dataset confirms that the lateral bodies are amorphous and reveal some connection sites linking them to the spikes of the palisade layer (Fig. 1b, Movie S2).

**Figure 1:**
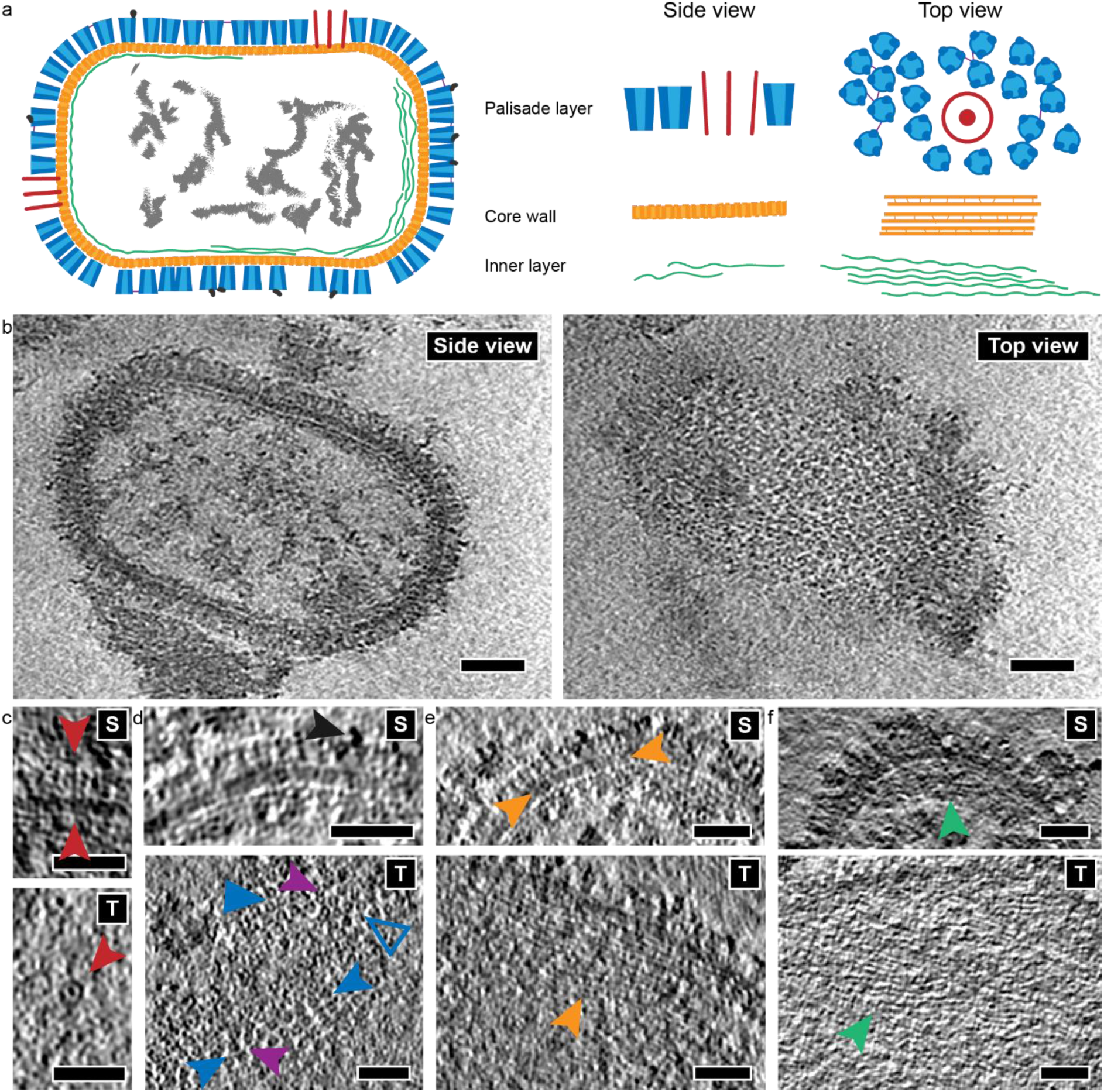
Appearance of the *in vitro* core. (a) Cartoon representation of the *in vitro* core based on the visual analysis of tomograms showing a slice through the middle of the core (left panel) and close ups for the three layers from the side and top view. The red structures, displayed as circles in the top view, represent the ring-like structures. (b) Representative example of side (S) and top (T) view of the *in vi*tro cores. The top layer displays the palisade of tube-like structures that have an irregular arrangement. The side view of the core reveals sparse density in the inner part of the core and the core wall consisting of three layers: the palisade (blue), the core wall (orange) and the most inner layer (green). (c-f) Close up of the three different layers surrounding the inner part of the core with the side and the top view. (c) A ring-like structure (red arrowhead). (d) Detail of the palisade layer at the surface of the core. The dark grey arrowhead points to the strong density occasionally observed on top of the palisade stakes. The blue filled triangle points to the hexagonal arrangement of the stakes while the hollow one to the similar arrangement, however consisting of eight, not seven stakes (including the central one). The blue arrowheads point to stakes arranged in rails, observed frequently. The violet arrowheads mark the thin threads connecting the stakes. (e) Detail of the core wall layer, which has stripe-like pattern as seen from the top (orange arrowheads). (f) Detail of the most inner layer from the side and top view (green arrowheads). Scale bars, 30 nm.

### AlphaFold2 predictions of major core proteins

AF2 [10] was used to predict the 3D structure of five viral proteins known to be abundant in fractions of cores prepared *in vitro* [4]. These are the gene products of A3, A4, A10, and L4 and the lateral body protein F17 [5]. The predictions were computed for various assemblies, from monomers to hexamers and different combinations of the proteins (for full list, see Table S1.1 to S1.3). Based on the confidence scores (Table S1.1), the monomers of A3, A10, and L4 were found to form ordered structures, while A4 and F17 are mostly disordered (Fig. S2a). The confidence scores for multimer predictions (Table S1.2) suggest that A3 and L4 could form dimers, while A10 is most likely to be a trimer.

### The palisade is composed of trimers of the major core protein A10

The initial low-resolution structure, obtained from the alignment of 400 manually picked stakes, displays a tube-like shape with a diameter of 9 nm and a length of 11.5 nm (Fig. S3c). From the AF2 predictions of the major core proteins, a trimer of A10 fits in shape and size to this first average (Fig. S3c). We used the AF2-predicted A10 trimer (Fig. 2a) as an initial reference to localize the stakes on all segmented cores using the oversampling approach (see Methods for more details). To limit potential template bias, the initial reference for the subsequent alignment was created using only the positions, not the orientations found during the localization (Fig. S2f). For the classification, we followed the protocol from [13]. First, we obtained 20 different *de novo* classes as potential candidates for further classification. Eight classes stand out with significant structural differences in the central parts of the stake (Fig S4a, b), while the remaining 12 classes are variations with minor differences in protomer positions and orientations, confirming the inherent flexibility of the stake structure. Following the classification protocol, the 8 classes were used as starting references for several independent classification runs (Fig. S4b). These revealed classes with distinct features within the central part of the stake. Specifically, they were characterized by the absence or presence of densities connecting the three protomers across the inner central region (Fig. S4b), and were assigned as hollow and connected trimers, respectively (Fig. 2b). Interestingly, mapping the classes back to the tomograms shows that there is no preferential clustering or location of the different classes on the core surface (Fig. S4d). The majority of classes form hollow trimers and have minor differences in orientation relative to the each other (Fig. S4b). The measured overall resolution of the best hollow and connected classes reached 7.7Å and 7.6Å, respectively.

**Figure 2:**
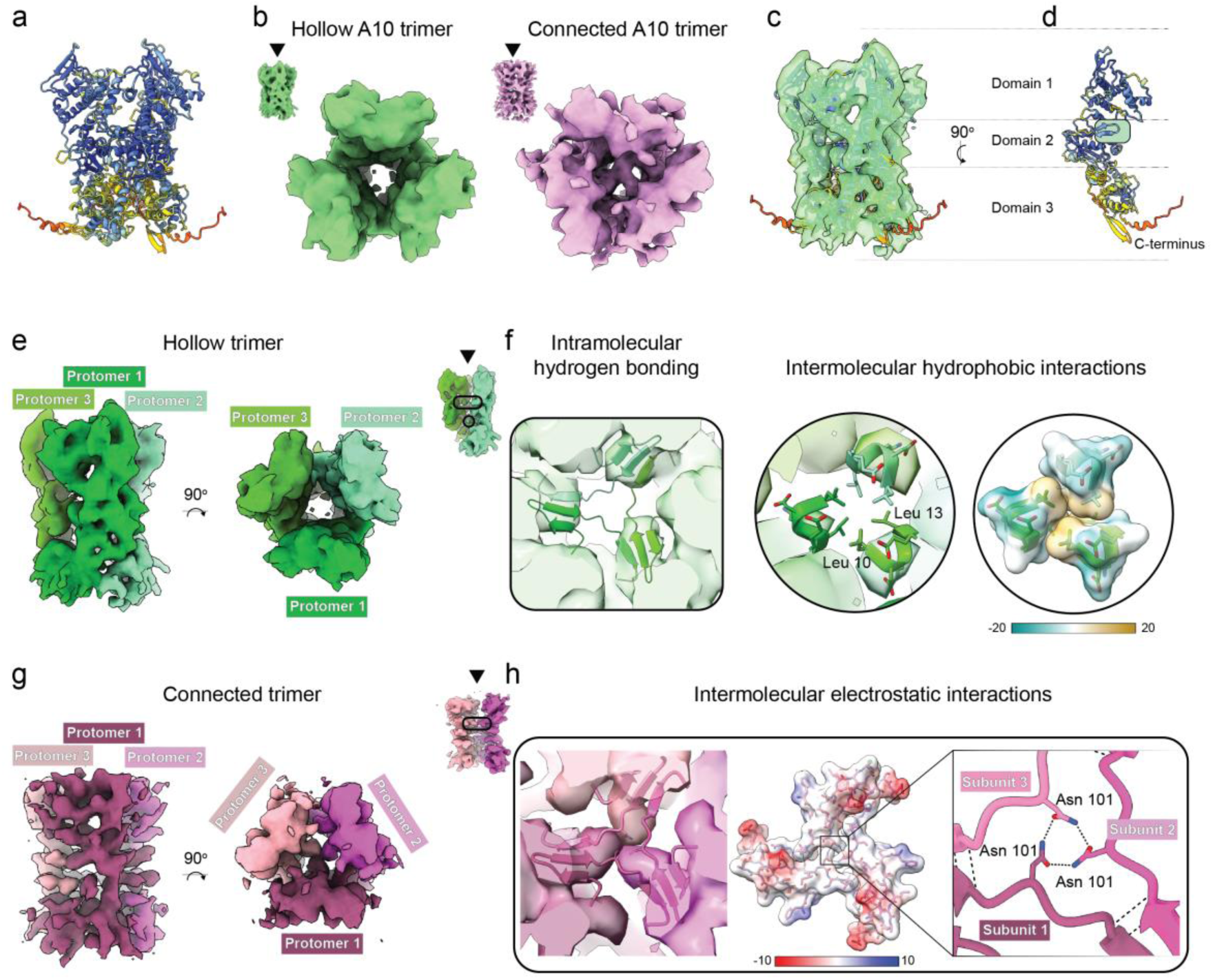
Classification and structural analysis of the palisade stalks made of A10 trimer. (a) AlphaFold2 prediction of A10 trimer colored by per-residue confidence score (pLDDT) between 0 and 100. Very high (pLDDT > 90, dark blue), confident (90 > pLDDT > 70, light blue), low (70 > pLDDT > 50, yellow), very low (pLDDT < 50, red). (b) Subtomogram averages (STA) of the two distinct A10 trimer classes, hollow (green) and connected (pink). (c) The AF2-predicted model of A10 trimer was rigid-body-fitted into the STA map of a hollow trimer. (d) The interface view of an individual protomer from the AF2-predicted A10 trimer. Coloring is same as in (a). The major interprotomer contact sites are highlighted with the green rectangle and green circle. (e) The STA density of the hollow trimer segmented into protomers colored in the shades of green. (f) The main contact points between protomers identified based on the fitting of AF2-predicted A10 trimer into STA density of the hollow trimer. The black rectangle shows the region for the interprotomer β sheets at the region of domain 2. The circles show the hydrophobic interface formed by the N-terminal residues Leu 10 and Leu 13 of each protomers. The hydrophobic surface is colored from dark cyan for the most hydrophylic to the dark gold for the most hydrophobic regions. The values correspond to the molecular lipophilicity potential calculated by the *fauchere* method of the mlp function in ChimeraX based on MLPP program. (g) The STA density of the connected trimer segmented into protomers colored in the shades of pink. (h) The main contact points between protomers identified based on the fitting of AF2-predicted A10 trimer into STA density of the connected trimer. The black rectangle shows the domain 2 region for the interprotomer beta sheets formed by two β strands of one protomer (Val 90 - Leu 92, and Asn 96 - Ile 99), and one β strand of the neighboring protomer (Ile 104 - Ser 107) is supported by electrostatic interactions of the Asn 101 residue of each protomer in the central channel of the trimer at the region of domain 2. The middle subpanel represents electrostatic potential of atoms calculated according to Coulomb’s law using Coulombic surface coloring tool in ChimeraX. The most negative potential colored in red, the most positive potential colored in blue.

Obtained STA structures indeed correspond to the trimers of A10 protein as suggested by the result of systematic fitting and subsequent molecular dynamics flexible fitting of the AF2-predicted model of A10 trimer into STA map of the hollow trimer (Fig S5a-d). This is supported by visual inspection of the secondary structures (Fig. S5e), and the model-to-map cross-correlation analysis (Fig. S5f). Notably, domain 3, which was predicted with the least confidence by AF2, also showed the lowest cross-correlation with the map (Fig. S5f). This, together with the high variability of domain 3 in STA structures, may suggest its role beyond forming the structural scaffold.

The scaffold of an A10 trimer and the major contact sites between its protomers were addressed by fitting of the AF2 model of the A10 trimer into the hollow and connected structures (Fig. 2e-h). While the hollow trimer is supported by a relatively weak hydrophobic interactions between Leu 10 and Leu 13 located near domain 3 (Fig. 2f, circular subpanel), in the connected trimer, protomers form three intermolecular beta sheets stabilized by electrostatic interactions between Asn 101 located in the flexible loop in the central channel of the trimer (Fig 2h).

### The palisade is an integral part of the incoming core during entry in cells

To exclude potential artifacts from the preparation of cores *in vitro*, we compare the structure of the palisade observed on cores *in vitro* and *in situ*. To this end, tomograms of cores found in the cytoplasm of host cells shortly after entry were acquired. In brief, HeLa cells grown on grids are plunge-frozen at 30 minutes post-infection without prior chemical fixation and thinned by focused ion beam (FIB)-milling (see Materials and Methods). Fifteen tilt series were acquired containing cytoplasmic cores and after tomogram reconstruction, 5 incoming cores were used for averaging of the palisade trimer as described above for the *in vitro* cores (Fig. 4a).

The overall architecture of the incoming cores observed *in situ* is very similar to the cores prepared *in vitro* (Fig. 1 and 3, Movie S3). The outer layer represents the prominent palisade (Fig. 3b) while the middle layer has a stripy appearance as highlighted above (Fig. 3a, c). The cytoplasmic cores are devoid of lateral bodies, known to dissociate immediately after entry and membrane fusion [5], [13]. The electron-dense layer found lining the core wall in the samples prepared *in vitro* is not seen in cytoplasmic cores (Fig. 3a). Instead, the central part of the core is homogeneously filled with filaments likely representing decondensed DNA (Fig. 3a), as described before [12]. A remarkable feature that was not observed *in vitro,* is the presence of strands attached to the palisade that have been previously reported by Hernandez et al. [9] and could be early transcripts leaving the core (Fig. 3d). Finally, the ring-like structures located between the palisade stakes found in *in vitro* cores (Fig. 1c) are also observed on the cytoplasmic cores (Fig. 3b).

**Figure 3:**
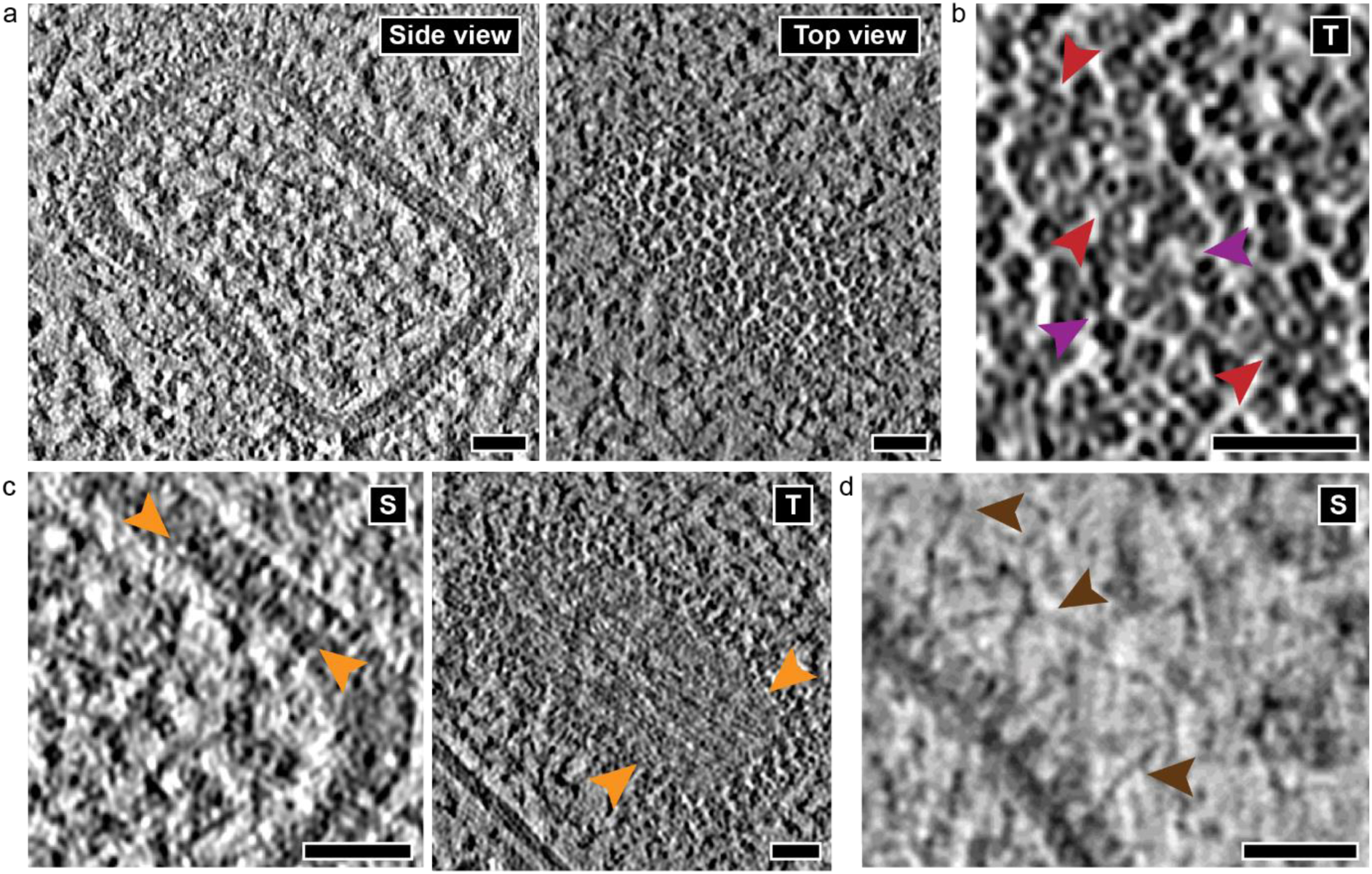
Analysis of incoming cores in the cytoplasm shortly after entry of VACV in HeLa cells. Slices through a representative tomogram from a core observed in the cytoplasm early in infection show a similar architecture as cores prepared *in vitro*. (a) Slice from a middle plane of the core (side view) and from the top (top view). The most inner layer observed on *in vitro* cores is missing and filaments without clear organization, potentially the viral dsDNA genome, can be observed. (b) Close-up of the top view from the palisade layer. Similar to *in vitro* cores, there is no regular arrangement of the stakes and the thin threads connecting the stakes can be observed (violet arrowheads). The red arrowheads point to the ring-like structures. (c) Side (S) and top (T) view of the core wall with the stripes indicated by the orange arrowheads. (d) Strands observed at the surface of the palisade (brown arrowheads). Scale bars, 30 nm.

### Comparison of the palisade spike *in vitro* and *in situ*

Despite the limited number of particles, due to challenges associated with the imaging of incoming cores in lamellae of infected cells, the structure of the palisade stakes from the *in situ* data was resolved to 14 Å. This is sufficient to conclude that the size and shape of the palisade from *in situ* cores is consistent with the *in vitro* one when compared at the same resolution (Fig. 4a). We further tested whether the stakes on intracellular cores are closer to the AF2 prediction model or our *in vitro* model by running alignment of the *in situ* data against both models. The constrained cross correlation scores were higher for the *in vitro*-based model than to the AF2 prediction of A10 trimer making us confident that the conformation of the stakes after classification of the subtomogram averages is physiologically relevant (Fig. 2). To further confirm the visual similarity between the *in vitro* and *in situ* cores, nearest-neighbor analysis was performed. The average distance between neighboring stakes is 8 nm for both datasets and the analysis of the normalized nearest neighbor position shows a similar trend (Fig. 4b, c). This analysis confirms the irregular arrangements of the lattice *in situ*, since there is no visible positional preference of the nearest neighbors. The number of neighboring stakes within a given distance per trimer shows that for both data sets only around 5% of the stakes form a hexagonal arrangement (Fig. 4c). The STA maps of pairs of trimers from *in vitro* cores show that even the neighboring trimers are not consistently oriented towards each other (Fig. 4d). The average number of stakes per core is 2690 for the *in vitro* data. Given that the *in situ* cores are not fully contained within the tomograms acquired on lamellae, we calculated an average stake density on the core surface by dividing the number of stakes per core by the surface area of the respective core segmentations. There is on average 0.018 stakes per 1 nm^2^ surface for *in situ* cores and 0.015 for *in vitro* cores.

**Figure 4:**
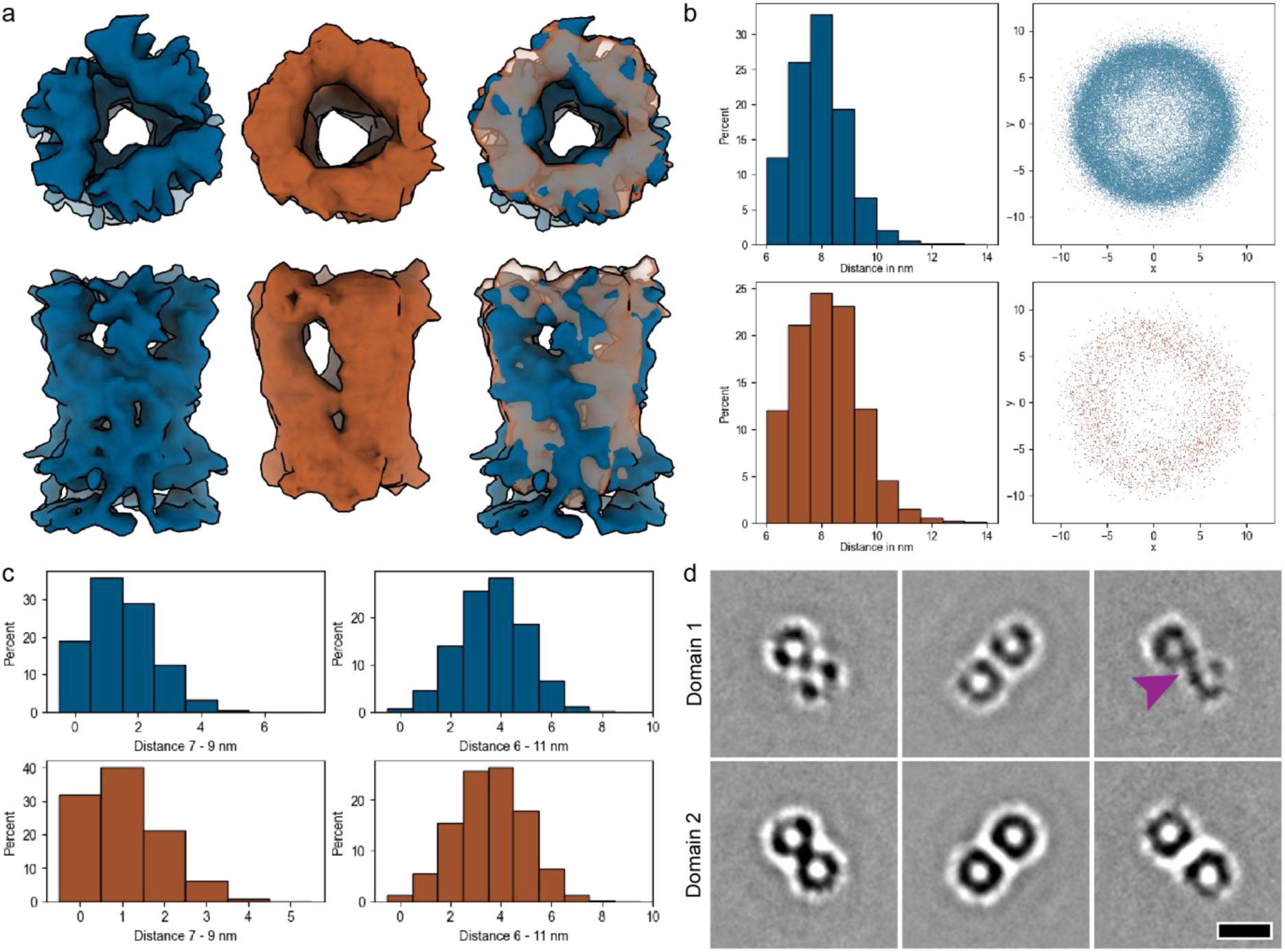
Comparison of *in vitro* and *in situ* palisade stakes. (a) The structures of *in vitro* (left, blue) and *in situ* (middle, brown) after localization and alignment (prior to classification). The right image shows their overlay. While the *in situ* map is less resolved (due to low number of particles and their large flexibility), the overall shape and size of both structures match very well. (b) Results of the nearest neighbor (NN) analysis performed on the trimers with top row being *in vitro* dataset and bottom row the *in situ* dataset. The histograms (right) show the average distances among the nearest neighbors, showing similar trend for both datasets. The relative position of the nearest neighbor to the normalized position and orientation of each stake is shown in the graphs on the right. The positions are shown in 2D for xy dimension, which corresponds to the top view of the map. Although minor clustering, corresponding to the hexagonal arrangement, can be observed in the *in vitro* dataset, the majority of stakes do not form any regular arrangements. (c) Distribution of number of NN with the distance constrained to the range of 7 nm to 9 nm (left) and the whole range, i.e. 6 nm to 11 nm (right). The constrained range corresponds to the average distance between stakes ±1 nm. At this distance the stakes have predominantly only 1 to 2 neighbors for both datasets. The whole range of distance is needed to obtained more NN and even in this case the majority of stakes has 3 to 4 NN. (d) STA maps of nearest neighbor pairs at distances 6.5 nm to 7.5 nm (left), 7.5 to 8.5 nm (middle) and 8.5 to 9.5 nm (right) from the *in vitro* dataset. The top row shows cross section from the top part of the domain 1, the bottom row is the cross section from the domain 2. Despite narrow distance ranges, the stakes do not have a form of a trimer as this shape was averaged out due to lack of strong preferred orientation among the pairs. At the distance from 8.5 to 9.5 nm a strong density connects the pair of stakes at the top of domain 1 (violet arrowhead) which corresponds to the observed threads from Fig 1. Scale bar 10 nm.

## Discussion

Among the first steps of the replicative cycle of VACV is fusion of the MV during entry and release of the viral core into the cytoplasm to initiate infection. This core has been studied to quite some extend, morphologically and biochemically, but the molecular composition of its sub-unit has remained elusive [5]. Cores can also be prepared from detergent-stripped virus, which facilitated previous studies [8] and our present analysis. By combining cryoET, STA and AF2, we show that the prominent palisade layer is composed of trimers of the gene product of A10. A10 is an abundant VACV core protein that is essential for the formation of the MV [14]; it is synthesized as a 110kDa precursor that is cleaved to a 65kDa form during virion maturation [15]. This maturation step is accompanied by the formation of the typical oval or brick-shaped core, displaying the palisade on its surface. As described before, the palisade units appear as trimers forming hollow tubes [8], [9]. Our study identified more than twenty different classes displaying minor differences in protomers orientations. Among those two global classes can be distinguished, one class presents an open, hollow, central part, whereas the other displays densities that connect the protomers across the center. This together confirms the structural flexibility that can be observed in tomograms where stakes appear to adopt slightly different shapes (Fig 1). The class with the best resolved features and highest resolution was used for model fitting with the AF2 prediction used as a starting model. It generally fitted the cryoET structure well, although the central channel of the trimer of the native structure, both *in vitro* and *in situ*, appeared to be more open. Most importantly, we show that the structure of the stakes on detergent-stripped virions is similar to the one on incoming core, making us confident that it is physiological relevant. In fact, the *in vitro* and *in situ* structures fitted each other better than the one predicted by AF2.

We found that the different classes are randomly distributed on the surface of the core without any clustering at specific locations such as the corners of the core. The stakes on both *in vitro* and *in situ* cores are randomly organized and display no specific pattern as shown by nearest neighbor analysis. This seems to contradict previous data, proposing that the palisade shows both unorganized regions as well as large patches with a hexagonal arrangement [3]. The difference can originate from the analysis of different stages of the virus cycle. The palisade may be more organized on the fully assembled infectious MV [3], [9] whereas it displays no obvious order on intermediates of disassembly, such as membrane-free cores.

The core is known to contain at least 5 abundant protein (see introduction), 4 of them are among the most abundant proteins of the MV, the gene products of A3, A10, A4 and F17 [16], implying important structural roles. F17, a 11kDa protein, is known to be part of the lateral bodies. These may remain attached to cores prepared by gentle detergent-stripping as in this study, but dissociates from the core immediately after entry [5]. AF2 predicts F17 to be generally unordered, as abundant component of the lateral bodies this fits their amorphous appearance. Our collective *in situ* and *in vitro* data allow us to draw the model of the molecular architecture as in Fig. 1a, where the palisade is composed of flexible A10-trimers. Attempts to localize the AF2 predictions of the other major core proteins within the tomograms were unsuccessful until now. Because these proteins are known to be part of viral core, the lack of high-confidence response within the inner core layer during attempted STA likely reflects discrepancies between the AF2 predictions and the experimental data at high resolution. The structure localization using oversampling surfaces with high-resolution map as initial reference is sensitive to the accuracy of the high-resolution information, especially in case of small structures or structures without distinct shape as is the case for the predictions for monomers of A3 and L4. The accompanying manuscript predicts that A3 constitutes the core wall below the palisade, a prediction that is consistent with immuno-labeling experiments on cytoplasmic cores [5]. Indeed, based on the AF2 prediction the monomer or dimer of A3 could fit size-wise into the core wall. In addition, the patch of strong negative charges on its surface could interact with the base of the A10 trimer which is positively charged. The location of A4, a 39kDa protein, remains unknown at this stage; previous immuno-labeling experiments on cytoplasmic cores would place this protein on the core surface. Indeed, our tomograms occasionally suggested the presence of an additional density on top of the palisade, but its characterization requires further analysis.

The identification of the molecular and structural composition of the palisade questions its function, elusive until this date. A10 is essential for the formation of the MV and its abundance suggests it to shape the core into its characteristic brick-shape and linking the core to the viral membrane and the lateral bodies. Indeed, we detected direct interactions between palisade units and the lateral bodies as described recently [9]. Accordingly, the palisade could fulfill a scaffolding function similar to the gene product of D13 during membrane assembly as recently suggested [9]. Being an abundant and flexible protein, we speculate the stakes to have additional functions related to the early replicative cycle, when the palisade is prominently exposed on the core surface. The cytoplasmic core is known to interact with microtubules [17] and the palisade stakes could mediate this interaction. The rings on the core surface, observed before [9] and in the accompanying study, are typically surrounded by A10-trimers. The latter might support, stabilize the rings or contribute to their typical ring-like structure. The subsequent step of the early infection is the uncoating of the DNA-genome and its release into the cytoplasm for replication. Images documenting this event suggest that genome-release occurs by a single and distinct break in the core wall [5], [12]. A distinct subset of A10 trimers could facilitate this opening, undergoing rearrangements that locally modifies the palisade layer and destabilizes the underlying core wall, for release of the viral DNA.

Altogether by combining cryoET, STA and AF2 we have identified the composition of the VACV palisade. The A10-trimer is an abundant and highly flexible protein, with the potential to fulfill multiple functions during the VACV assembly and disassembly, the focus of future research.

## Acknowledgments

We thank Stefanie Boehm (MPI of Biophysics) for critical reading of the manuscript and helpful discussion. We thank Sylvia Panitz and Regina Eberle, microscopy facility Paul Ehrlich Institute, for excellent technical assistance. We also acknowledge support by Cornelia Cazey, Ulrike Laugks, and Carolin Seuring, for access to the Cryo-EM multi-user facility at CSSB and providing time for sample preparation, screening, and data collection; Wolfgang Lugmayr for help and support in using the CSSB partition on the DESY computer cluster for cryo-EM data processing. Part of this work was performed at the Cryo-EM multi-user Facility at CSSB, headed by K.G. and supported by the UHH and DFG (grants INST 152/772-1, 774-1, 775-1 and 776-1). In the framework of this project, E.R.J.Q. was supported by an individual fellowship from the Alexander von Humboldt Foundation and a Klaus Tschira Boost Fund. JKL is supported by the excellence cluster Loewe DRUID of the state Hesse, project E7-P. JL is supported by DFG (project 515013236).

## Contributions

JL, SCD, VP, DV, ERJQ, and BT performed data processing and subtomogram averaging. SCD, KB, SW, SRT, and ERJQ optimized sample preparation and performed data collection. JL, SCD, and AOK performed analysis on AF2 models. JL, IK, AOK did the model fitting. KG, ERJQ, BT, JKL designed the experiments. JL, SCD, ERJQ, BT and JKL wrote the manuscript while VP, DV and KG did a critical manuscript review.

## Data availability

The electron microscopy density maps of the A10 trimers have been deposited in the Electron Microscopy Data Bank under accession codes: EMD-XXXX, EMD-YYYY, EMD-ZZZZ (loose and tight conformation from the *in vitro* dataset, trimer from the *in situ* data). The raw tilt series of the in vitro dataset together with representative tomograms from both *in vitro* and *in situ* datasets were deposited at EMPIAR-QQQQ. The python scripts developed for core picking and contextual analysis will be publicly available on github at the time of the publication.

## Materials and Methods

### Mature virions

#### Sample preparation

Mature virions (MV) of VACV strain Western Reserve (WR) were purified as previously described [8]. The preparation was sonicated 2×30 seconds in a waterbath sonicator prior to be mixed in a 1:1 ratio with 5 nm gold fiducials (UMC Utrecht). This solution was then applied to Quantifoil™ 200 mesh R2/2 copper grids prior to plunge freezing into liquid ethane/propane cooled to liquid nitrogen temperature using a Vitrobot Mark4 (Thermo Scientific) after blotting for 3 seconds at 4°C.

#### Data collection

At the Cryo-EM multi-user facility at CSSB, the grids were imaged using a Titan Krios G3 operated at 300 kV (Thermo Scientific) and equipped with a BioQuantum-K3 imaging filter (Gatan). Grids were screened and tilt series acquired using SerialEM [18]. The low-dose mode was setup to achieve a dose of ~15 e-/pixel/sec on the detector directly on the sample with a 70 μm objective aperture and a 20 eV energy slit width. Tilt series were collected with the dose-symmetric tilt scheme [18] over a +/-60 degrees tilt range with a 3 degree increment step. Movie frames were recorded in electron counting mode with a pixel size of 1.372 Å, a target defocus of −3 µm and a total dose of 1.5 e-/Å^2^ over 10 frames, resulting in a total dose of ~80 e-/Å^2^ for each tilt series.

#### Data processing

Tomograms were processed within eTomo (IMOD 4.11 package) [19]. Prior to the reconstruction, the tilt series were aligned using the fiducial-based alignment and dose-weighting was applied to remove the damaging frequencies. The tomogram reconstruction was done with 3D-CTF correction and SIRT-like filter equivalent to 7 iterations.

### Cryo electron tomography of viral cores prepared *in vitro*

#### Sample preparation

Purified VACV MVs were diluted in 50 mM TrisCl, pH 8.5 and mixed with an equal volume of 0.2% Nonidet P-40 and 20 mM DTT diluted in Tris-CL. The mixture was sonicated 3 times for 1 minute in a waterbath sonicator and incubated in a ThermoMixer (Eppendorf) for 30 minutes at 37°C and 500 rpm. The mixture was layered on a 36% (w/v) sucrose in 50 mM Tris, pH 8.5 and the cores pelleted in a table top centrifuge at 100,000 rpm. The sucrose was removed without disturbing the translucent pellet and remaining sucrose drained by placing the tube top down for several minutes on Whatman paper. The pellet was resuspended in 50 mM Tris-Cl and placed on ice. For cryo electron tomography, freshly prepared cores were water bath-sonicated for 3×1 min and mixed 4:1 (cores:gold) with 10 nm gold particles coupled to protein A (Cell Microscopy Core, Utrecht University). Cu/C, R2/2, 200 mesh grids (Quantifoil) were glow-discharged for 90 sec using a Pelco easiGlow device. 3μl viral core sample were applied to freshly glow discharged grids and grids were plunge frozen in liquid ethane using a Vitrobot Mark4 (Thermo Scientific) at 4°C and 100% relative humidity. Vitrified grids were stored in liquid nitrogen until further use.

#### Data collection

Tilt series data of purified cores were acquired using SerialEM software [19] on a Titan Krios G4 transmission electron microscope (Thermo Scientific), operated at 300kV and equipped with e-CFEG, SelectrisX energy filter and Falcon4 direct electron detector, essentially as described in [20]. Grids were initially mapped at low magnification in 6×6 montages. Intermediate magnification maps of suitable grid squares were collected with a bin 1 pixel size of 19.4 Å, −100 μm defocus, a 70 μm objective aperture and a 20 eV energy slit. SerialEM low-dose mode was set up to achieve ~7 e −/pixel/sec camera dose-rate with a 70 μm objective aperture and a 20 eV energy slit inserted. Tilt series data were collected in a dose-symmetric tilt scheme [18] over a +/-60 degrees tilt range with 3 degrees tilt increments in groups of 2 tilts. Tilt images were acquired in electron counting mode with a pixel size of 1.577 Å, a target defocus of −1.5 to −4.5 µm and a total dose of 3.0 to 3.1 e-/Å2 over ten frames, resulting in a total dose of ~125 e-/Å2 per tilt series.

### Cryo electron tomography of viral cores *in situ*

#### Sample preparation

To image cores *in situ*, HeLa cells were grown overnight on Au/Au, R1.2/1.3, 200 mesh grids (Quantifoil) at 37°C and 5% CO_2_. Cells were placed in serum-free DMEM high-glucose medium (Gibco) and virus (WR strain with A3 protein tagged with YFP, kindly provided to us by Michael Way, The Francis Crick Institute, UK) bound for 45 minutes at RT followed by 30 minutes at 37^◦^C for virus entry. Grids were plunge-frozen in liquid ethane using EM GP2 device (Leica Microsystems). Lamellae were prepared on an Aquilos 2 (Thermo Scientific), operated by Maps and AutoTEM software (Thermo Scientific). Sample preparation, milling and polishing steps were done automatically using AutoTEM with a milling angle target of 8 degrees (2 degrees of tolerance) and a final lamella thickness set to 120 nm.

#### Data collection

For tilt series acquisition of *in situ* cores in lamellae of infected HeLa cells, grids were imaged using a Titan Krios G3 operated at 300 kV (Thermo Scientific), equipped with a BioQuantum-K3 imaging filter (Gatan). Grids were screened and data acquired using SerialEM. First, grids were mapped at low magnification to localize the lamellae. Mapped grids were pre-tilted to the angle used during milling (specific for each lamella between 7 and 10 degrees), which was the starting angle for tilt series acquisition. SerialEM low-dose mode was setup to achieve ~15 e /-pixel/sec camera on lamella with a 70 μm objective aperture and a 20 eV energy slit inserted. Tilt series data were collected in a dose-symmetric tilt scheme [18] over a +/-60 degrees tilt range around the starting angle with a 3 degrees tilt increment. Tilt images were acquired in electron counting mode with a pixel size of 2.653 Å, a target defocus of −5µm and a total dose of 4.5 e-/Å2 over ten frames, resulting in a total dose of ~200 e-/Å2 per tilt series.

### Preprocessing and tomogram reconstruction

The defocus for each tilt series was estimated with CTFFind4 [21]. The tilt series were dose-weighted using the MATLAB script from [22]. Suboptimal projections or whole tilt series were removed from dataset, resulting in 34 tilt series. They were aligned using fiducial-based alignment within eTomo. The tomograms were reconstructed from 8x binned tilt-series using WBP with SIRT-like filter option. These tomograms were used only for picking of cores within Napari (see below). The unbinned tomograms were reconstructed using novaCTF [23] with phaseflip correction, astigmatism correction, and a 15nm slab size. These tomograms were binned 2x, 4x, and 8x using Fourier3D [24]. These 3D-CTF-corrected tomograms were used for subtomogram averaging (STA). For the complete parameter setup of individual steps see Table S2.

### Subtomogram averaging (*in vitro*)

#### Created initial positions on the core surface

The cores were manually segmented by drawing contours around the cores in multiple layers of 8x binned tomograms in Napari. The layers were used to create the complete surface of the virions using the triangulation of the respective convex hulls. The resulting surfaces were then used to determine the starting position and orientations for STA (Fig S3a,b). Two sets of positions were produced for different purposes. The first set was generated for STA with an initial reference. For this set the surface was sampled at distance of 3.7 nm. The second set of positions was created specifically for resetting normal vector in *de novo* STA (see below). Surface for this set was sampled with distance of 1.2 nm.

#### Initial alignment

A total of 400 stakes were manually picked to generate the initial reference in novaSTA (Table S2). Subsequently, STA using all subtomograms was done in novaSTA [25] on 8x binned tomograms (Table S2). After the alignment, overlapping subtomograms were removed from further processing together with the subtomograms located near the carbon edge. The visual inspection of the aligned positions showed that the stakes are only partially covered and many subtomograms were aligned into the empty regions among them as these regions often resembled shape- and size-wise the stakes at this resolution. High cross-correlation threshold was used to select only subtomograms that were correctly located and obtained initial STA map from ~8000 subtomograms which gave us an estimate of the size and shape of the stakes (Fig S3c).

#### AlphaFold2 prediction

AlphaFold-Monomer version 2.2.0 [10] was used to generate models of four monomers of proteins A3, A4, A10, and L4 (Fig S2). AlphaFold-Multimer version 2.2.0 [26] was used to create multimers of these proteins as well as their combinations (see Table S1 for the full list). The AF2 parameters were set to default, except for the max_recycles parameter which was set to 12, to ensure convergence of the modeling. The models of the monomers were scored according to the predicted pLDDT score, while the models of protein complexes were scored using a combined ipTM (interface predicted TM-score) and pTM score (predicted TM-score), as returned by AF2 (Table S2).

#### *De novo* subtomogram averaging

The overall shape of the initial STA map matched the AF2 prediction model of a trimer of A10 (Fig S3c). The AF2 model of the trimer was used as initial reference for STA on the oversampled surfaces. However, to utilize the high-resolution necessary to distinguish between the stakes and the regions among them the STA was done on 2x binned tomograms. The alignment resulted in correctly placed positions (Fig S3d) and a map resembling A10 trimer (Fig S3e). To avoid template bias, the orientations found during the STA were neglected. Instead, each subtomogram was assigned orientation based on its position on the surface. More precisely, the closest point from the second oversampled set was found and its normal vector was used to assign the cone angle of the subtomogram. The in-plane angle was assigned randomly. An STA average produced with the new orientation resembled featureless tube (Fig 3f). This was used as a new starting reference for the STA that was done using STOPGAP [27]. The overview of the full parameter setup is in Table S2. The number of particles contributing to the STA map was 114299.

#### Classification

The visual inspection of tomograms suggested high flexibility of the stakes. The protocol from [20] was used to perform classification on the stakes using STOPGAP. In total 20 *de novo* initial references were created by choosing random subsets of particles (10 subset contained 10 000 particles, 10 subsets 5000 particles). Two independent STA runs of 15 iterations were done using simulated annealing for first 10 iterations (with temperature factor 10) and random search for the last 5 iterations. All twenty classes contained a trimer, however 8 classes contained distinct features separating them from others (Fig 4a). These 8 classes were used as initial references for 4 independent STA runs with random search. In total 10 iterations were enough for all the runs to stabilize (Fig 4c). Subsequently, the consistency cleaning was performed; only particles that ended in the same class for all 4 runs were kept, the rest was discarded. The remaining particles underwent 7 iterations of STA multiclass alignment (i.e., without the possibility to change the class). Class 6 (reported resolution 7.6Å, 14915 particles) and class 7 (reported resolution 7.7Å, 5161 particles) were used for further analysis as the tight trimer and the loose trimer, respectively. The 8 classes were mapped back to tomograms using ArtiaX plugin [28] for ChimeraX [29] to inspect possible clustering of classes. The model created using ArtiaX shows that there is no connection between the subtomogram class and its position on the cores.

### Subtomogram averaging (*in situ*)

The STA on *in situ* data was performed in the same way as for the *in vitro* dataset with the exception of the initial alignment, which was skipped. Instead, after segmenting the cores, the STA was directly performed with AF2 model of A10 trimer. After the localization, the new orientations were assigned, followed by STA using STOPGAP. The trimer was resolved to the resolution of 13.4Å (Fig 4). The relatively low number of particles (6325) prevented any attempts for classification. Table S2 contains all the parameters for the *in situ* STA.

### Systematic fitting of AlphaFold models to cryo-EM maps

We followed the procedure from the work of Zimmerli et al. [30] to locate the AlphaFold models in cryo-EM map. The flexible chain at the N-terminus (Met 1 - Thr 9) and at the C-terminus (Ser 592 - Gly 614) of the model were removed due to no extra densities observed in that region. All the models were filtered to 9 Å, and the resulting simulated model maps were subsequently fitted into cryo-EM maps by global fitting as implemented in UCSF Chimera [31] using scripts within Assembline [32]. The fitting was perform using 10,000 random initial placements, with at least 30% of the model map covered by the cryo-EM map. To evaluate the fit of each model, we used the cross-correlation about the mean (cam score), which was calculated using UCSF Chimera [31]. We assessed the statistical significance of each fitted model using a p-value derived from the cam scores. To obtain the p-values, we first transformed the cross-correlation scores to z-scores using Fisher’s z-transform, computed two-sided p-values based on an empirical null distribution derived from all non-redundant fits, and corrected the p-values for multiple testing using the Benjamini-Hochberg procedure [33].

### Flexible fitting and model analysis

To improve the fit of the A10 trimer, the molecular dynamics flexible fitting was performed using the model obtained by systematic fitting and the cryoET density map of the hollow trimer as inputs. The flexible fitting was performed in the web server of the Namdinator pipeline tool [34] in two sequential cycles with the following parameters: map resolution 12 Å, start temperature 500 K, final temperature 298 K, G-force scaling factor 0.3, minimization steps 5000, simulation steps 20000, phenix real space refinement cycles 0, implicit solvent excluded. The obtained model was subjected to phenix cryo-EM validation tool [35] to calculate model-to-map cross-correlation and the model-map Fourier shell correlation using default parameters and resolution set to 10 Å. The model to map fitting cross-correlation was calculated for the coordinates of one protomer and its corresponding density, as well as for individual domains of the protomer and their respective densities. All densities were segmented in ChimeraX [29] using the segger function [36] based on visual inspection.

For the analysis of the interprotomer contacts, specifically hydrophobic interactions between the alpha helixes Leu 10 – Leu 13 of domain 3 (Fig. 2f), and electrostatic interaction of the beta sheets Val 90 - Leu 92, Asn 96 - Ile 99, and Ile 104 - Ser 107 of domain 3 (Fig. 2h), the AF-predicted model was used. First, the trimer was fitted as a rigid body, then the corresponding regions were selected separately, and fitted as two individual rigid bodies into the densities of hollow and connected trimers. The molecular lipophilicity potential was calculated using the default fauchere method of the mlp function in ChimeraX based on MLPP program [37]. The electrostatic interactions between were calculated in ChimeraX of the Asn 101 residue of each protomer in the central channel of the trimer at the region of domain 2. The electrostatic potential of the beta-sheet-forming residues at the domain 2 region was calculated according to Coulomb’s law using the Coulombic surface coloring tool in ChimeraX.

---

**Supplementary Table 1.1.** List of the proteins of VACV analyzed in this study. The average per-residue confidence score (pLDDT) is given for the highest ranked model for the different proteins, along with their corresponding Uniprot IDs. The pLDDT is the confidence measure of AlphaFold2. The score ranges from 100 (indicating perfect confidence) to 0 (indicating no confidence).

| protein | Uniprot ID | pLDDTs |
| --- | --- | --- |
| **A3** | P06440 | 79.7 |
| **A4** | P29191 | 51.7 |
| **A10** | P16715 | 86.1 |
| **L4** | P03295 | 90.1 |
| **f17** | P07396 | 71.8 |

**Supplementary Table 1.2.** List of the model score from the AlphaFold multimer software for the highest ranked model of proteins listed in Table S1.1 when different symmetries are applied. The score represents the confidence of the protein-protein interface prediction. It is calculated using the Interface pTM (iptm) score and pTM score. The score ranges from 1.0 (indicating perfect confidence) to 0 (indicating no confidence).

| protein | oligomer | iptm+ptm |
| --- | --- | --- |
| **A4** | dimer | 0.224 |
|  | trimer | 0.283 |
|  | tetramer | 0.241 |
|  | pentamer | 0.175 |
|  | hexamer | 0.188 |
| **A3** | dimer | 0.761 |
|  | trimer | 0.289 |
|  | tetramer | 0.414 |
|  | pentamer | 0.355 |
| **A10** | dimer | 0.244 |
|  | trimer | 0.823 |
| **L4** | dimer | 0.700 |
|  | trimer | 0.163 |
|  | tetramer | 0.152 |
|  | pentamer | 0.167 |
|  | hexamer | 0.167 |
| **f17** | dimer | 0.266 |
|  | trimer | 0.186 |

**Supplementary Table 1.3.**
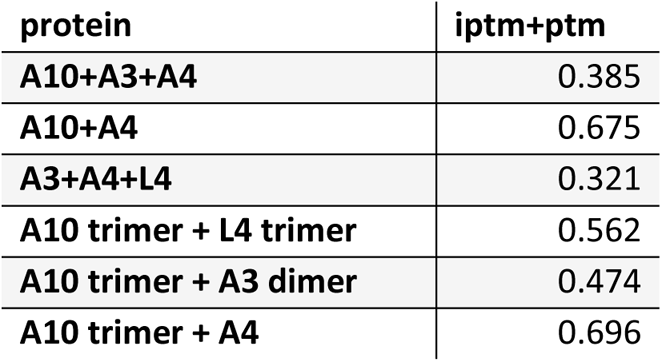
List of the model score from the AlphaFold 2 multimer software for proteins listed in Table S1.1 in different combinations.

**Supplementary Table 2.**
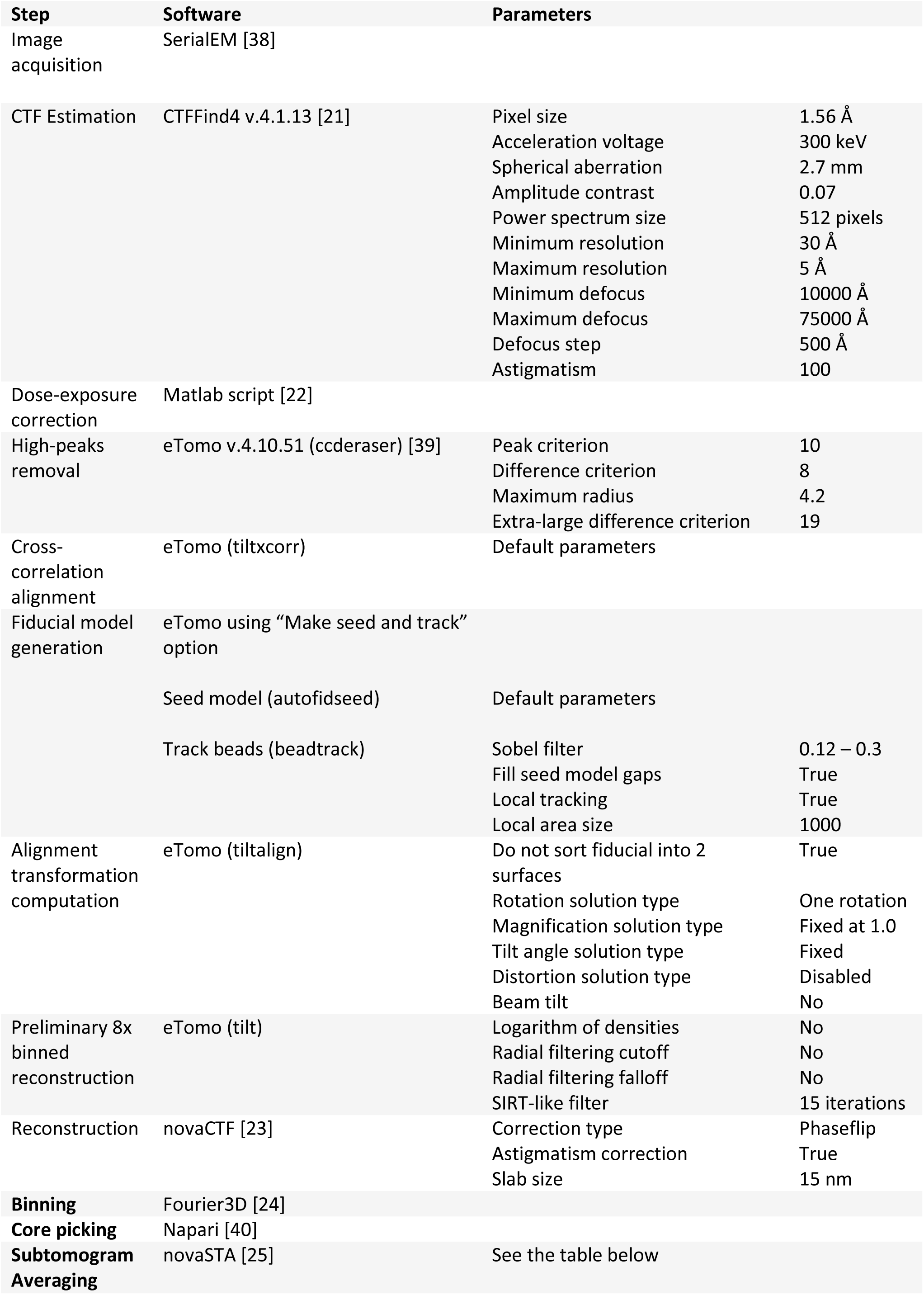
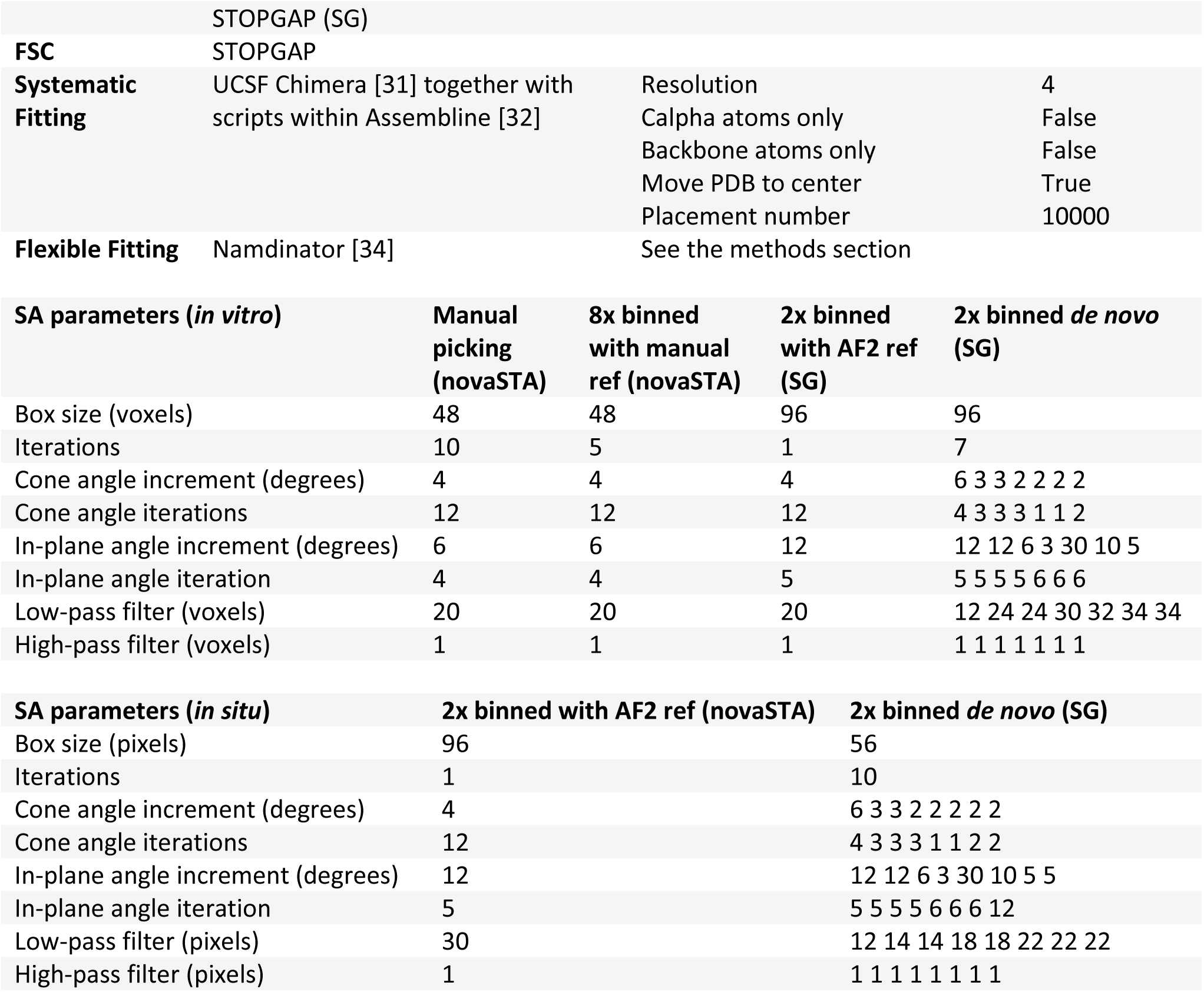
Overview of software and parameters used for processing.

**Supplementary Figure 1:**
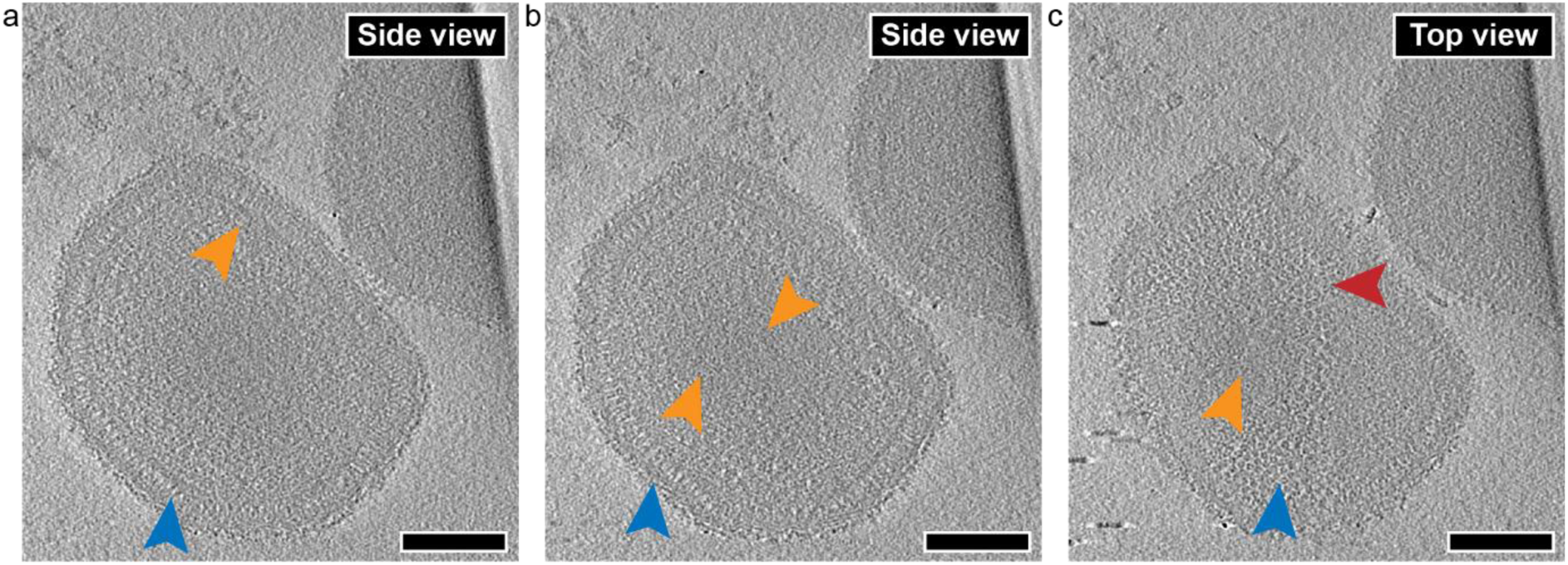
Slices of a tomogram of purified mature virion (MV) of VACV. (a) Slice of a tomogram (Movie S1) showing a cross-section of a MV with the core wall (orange arrowhead) underneath the viral envelope. Yellow arrowheads show side views of the palisade layer. (b) slice with the middle layer appearing as stripes (orange arrowheads). (c) slice closer to the top of the MV showing a top view of the palisade and the ring-like structures (red circle). Scale bars, 100 nm.

**Supplementary Figure 2:**
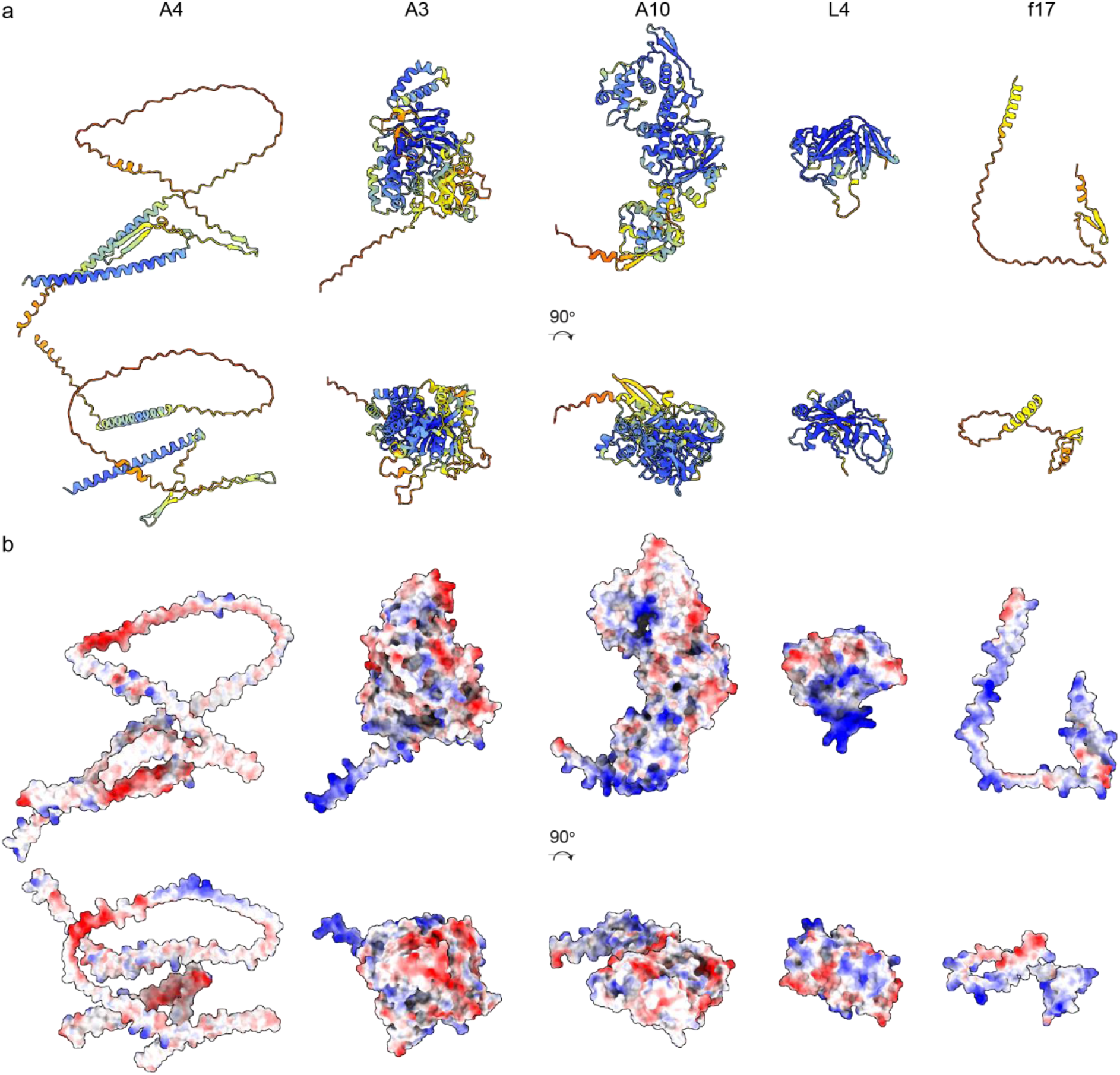
Structures of monomers predicted with AlphaFold2 software. The best model predicted for A4, A3, A10, L4 and F17, is depicted (a) colored according to plddts scores. (b) electrostatic surface potentials are colored red and blue for negative and positive charges, respectively, and white color represents neutral residues.

**Supplementary Figure 3:**
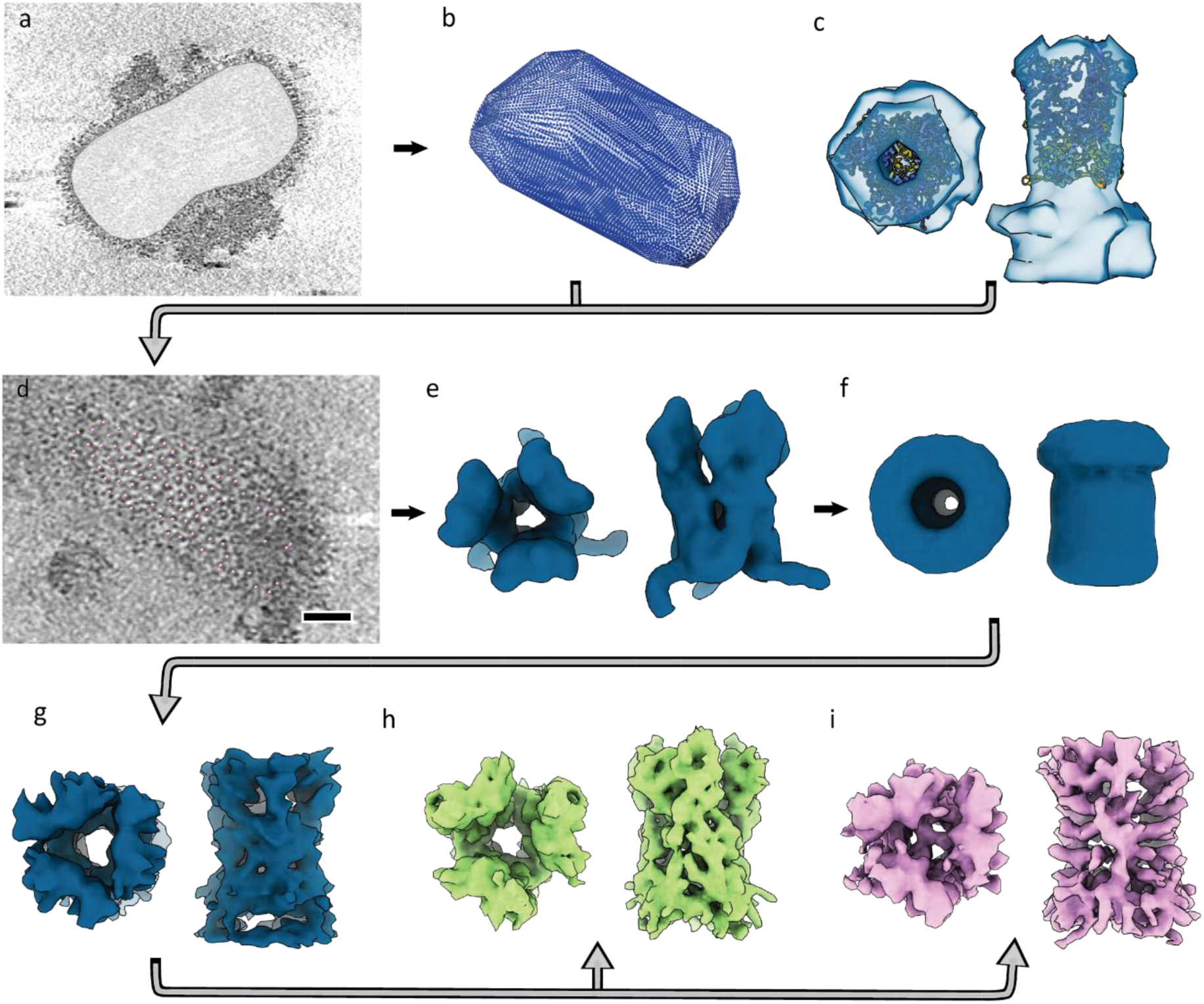
Particle picking and subtomogram averaging workflow. (a) Contour layer of the core in Napari. (b) Oversampled positions on the core surface. (c) The map generated by subtomogram averaging (STA) after manual picking of the stakes with the AlphaFold2 (AF2) prediction model fitted inside using Chimera. (d) The locking of the subtomograms onto the correct positions using the AF2 prediction model as the initial reference. (e) The STA map generated using the AF2 prediction model as the reference. (f) The STA map obtained after assigning new orientations based on their position on the core surface. (g) The *de novo* STA map generated from the new surface-based orientations. (h) Loose A10 trimer produced after the classification in STOPGAP. (i) Connected A10 trimer after the classification in STOPGAP.

**Supplementary Figure 4:**
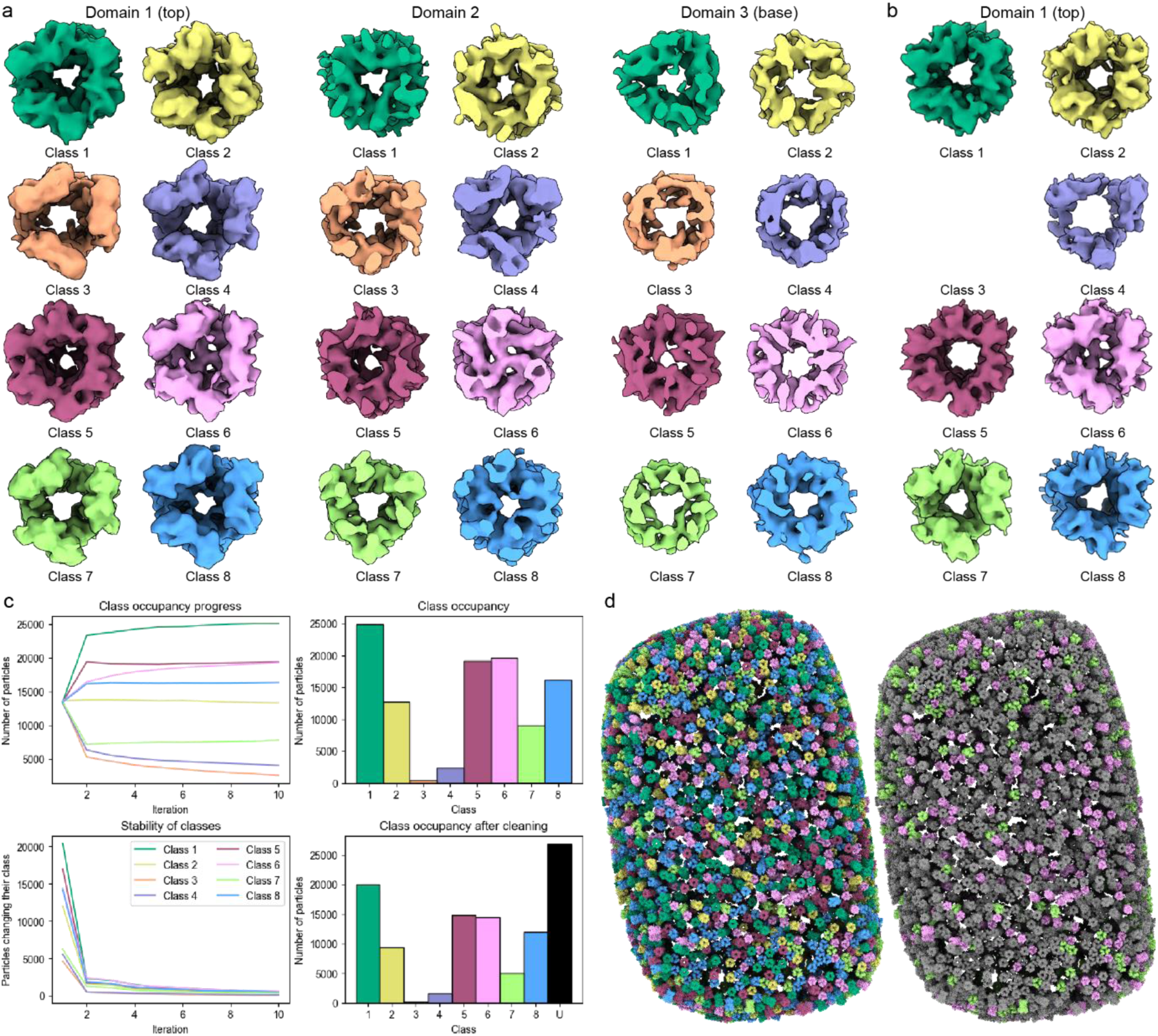
Classification of A10 trimer. (a) The classification with the starting references that were created by random subsets of particles yielded 8 different classes. The differences were mostly in the additional densities in the domain 2 (class 2, 4, 6) and domain 3 (class 1, 3, 5, 8). (b) The final 8 classes produced by 4 independent STA runs with the classes from (a) used as starting references. (c) Graphs showing class occupancy progress (top) and particle stability (bottom left) from a single STA run with the starting references from (a). Four independent and random STA runs were done and only particles that ended up in the same class for all 4 runs were kept (bottom right). The rest remained unassigned (U). (d) The final maps were plotted back to tomograms using ArtiaX plugin in ChimeraX. The color-coding corresponds to the individual classes (left panel) or classes 6 (tight trimer) and 7 only (loose trimer) (right panel). There is no apparent clustering or preferred position on the core palisade that would correspond to the assigned classes.

**Supplementary Figure 5:**
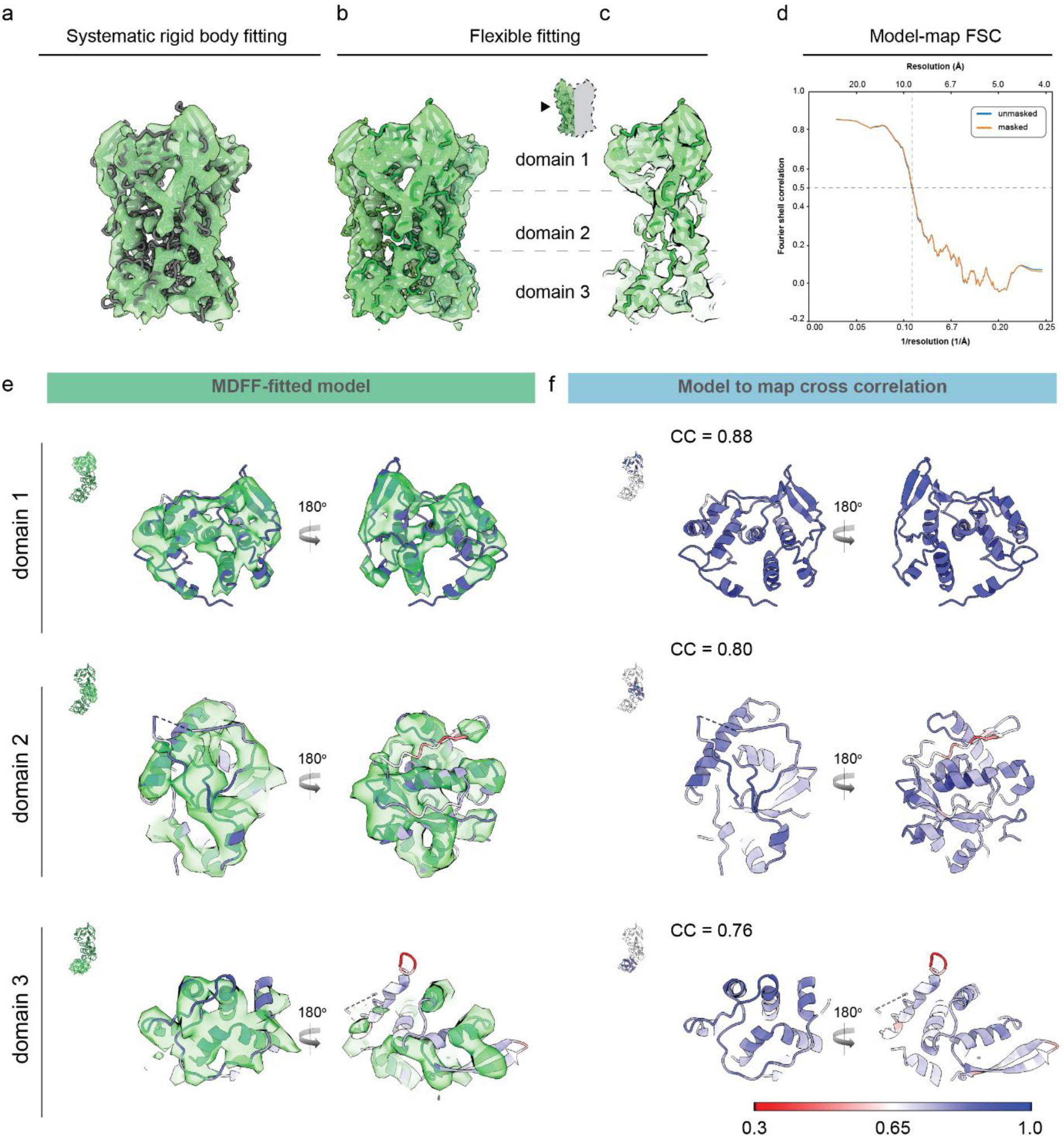
A10 trimer model fitting and validation. (a) The model of an AF-predicted A10 trimer (dark grey) after systematic rigid body fitting into the cryoET density of a hollow trimer (green). (b) The model of an A10 trimer (dark green) after molecular dynamics flexible fitting (MDFF) into the cryoET density of a hollow trimer (green). The model from panel (a) was used as an initial template. (c) Clipped view of panel (b) showing the model fit of one protomer. (d) The model-map Fourier shell correlation (FSC) calculated for the model of one protomer shown in panel (c). (e) Per domain representation of the protomer fitting into cryoET density of a hollow trimer. Solvent view is shown on the left, interface view is shown on the right. (f) Per domain representation of the protomer (same as in (e) with residues colored according to the model-to-map cross-correlation (CC) scores calculated in phenix cryoEM validation tool, from the lowest (red) to highest (blue). Solvent view is shown on the left, interface view is shown on the right. The average CC scores for each domain is indicated in the upper left corner.

Supplementary movie 1: Tomogram of MV core.

Supplementary movie 2: Tomogram of two *in vitro* cores.

Supplementary movie 3: Tomogram of *in situ* core.

